# Dissecting Mismatch Negativity: Early and Late Subcomponents for Detecting Deviants in Local and Global Sequence Regularities

**DOI:** 10.1101/2023.11.20.567982

**Authors:** Yiyuan Teresa Huang, Chien-Te Wu, Shinsuke Koike, Zenas C. Chao

**Affiliations:** International Research Center for Neurointelligence (WPI-IRCN), UTIAS, The University of Tokyo, 113-0033, Japan; School of Occupational Therapy, College of Medicine, National Taiwan University, Taipei, Taiwan; Department of Multidisciplinary Sciences, Graduate School of Arts and Sciences, The University of Tokyo, Tokyo, Japan; University of Tokyo Institute for Diversity & Adaptation of Human Mind (UTIDAHM), Tokyo, Japan

**Keywords:** Mismatch negativity, hierarchy, subcomponents, predictive coding, EEG

## Abstract

Mismatch negativity (MMN) is commonly recognized as a neural signal of prediction error evoked by deviants in the expected pattern of sensory input. Studies show that MMN diminishes when a sequence pattern becomes more predictable over a longer timescale. This implies that MMN is comprised of multiple subcomponents, each responding to different levels of temporal regularities. To probe the hypothesized subcomponents in MMN, we record human electroencephalography during an auditory local-global oddball paradigm where the tone-to-tone transition probability (local regularity) and the overall sequence probability (global regularity) are manipulated to control temporal predictabilities at two hierarchical levels. We find that the size of MMN is correlated with both probabilities and the spatiotemporal structure of MMN can be decomposed into two distinct subcomponents. Both subcomponents appear as negative waveforms which peak early in the central-frontal area and late in a more frontal area, respectively. With a quantitative predictive coding model, we map the early and late subcomponents to the prediction errors that are tied to local and global regularities, respectively. Our study highlights the hierarchical complexity of MMN and offers an experimental and analytical platform for developing a multi-tiered neural marker, applicable in clinical settings.

## 1. Introduction

Mismatch negativity (MMN) is a difference in event-related potentials between unexpected (deviant) and expected (standard) sensory events, occurring at 100-250 msec after event onset and over the frontal-central brain area, (Näätänen et al., 2007; Sams et al., 1984). It is a key biomarker to evaluate the capacity for statistical learning and deviance detection in individuals with autism spectrum disorder (ASD) (Dunn et al., 2008), schizophrenia (Erickson et al., 2016; Fisher et al., 2012; Koshiyama et al., 2020), and developmental delay (Kujala & Näätänen, 2001).

MMN has been thought to represent a prediction error (Garrido et al., 2009; Wacongne et al., 2012). Its characteristic amplitude and latency are influenced not only by bottom-up sensory inputs, e.g., a more pronounced pitch difference between standard and deviant auditory stimuli yielding a larger MMN with an earlier peak (Pakarinen et al., 2007), but also by top-down predictions established through learning, e.g., less predictable stimuli typically evoking a larger MMN with an earlier peak (Javitt et al., 1998; Sabri & Campbell, 2001; Sato et al., 2000). Studies have also shown that when dealing with stimuli containing multi-layered sequential sequence structures, a smaller MMN response is triggered when the stimulus becomes more predictable as the temporal regularity over a longer timescale is considered. This predictability can be demonstrated through sequences that adhere to descending or ascending tonal order (Ruhnau et al., 2012; Wiens et al., 2019) or a deviant stimulus often occurring after certain numbers of standard stimuli (Goris et al., 2018; Wacongne et al., 2011; Yaron et al., 2012). This evidence indicates that MMN is not a unitary signature but a composite of multiple subcomponents, each corresponding to different levels of temporal statistics and degrees of prediction errors. However, the distinct neural signatures of these hypothesized subcomponents, as well as their individual contributions to the overall structure of MMN, remain unclear.

Here, we record human electroencephalography (EEG), while participants are listening to a local-global oddball paradigm (Bekinschtein et al., 2009) which comprises local and global temporal regularities in tone sequences. To dissociate MMN subcomponents representing the hierarchical regularities, we manipulate both the tone-to-tone transition probabilities at the local level and sequence probabilities at the global level to create various sequence blocks. We then analyze the EEG responses across these different blocks using an unbiased data-driven decomposition approach (Chao et al., 2018). Two subcomponents, each with unique spatiotemporal signatures, are extract from the observed MMN and provide the best explanation for the data. The activation patterns of these two subcomponents, in alignment with a hierarchical predictive coding model (Chao et al., 2022), suggest that the early and late subcomponents represent the local and global prediction-error signals. Our findings reveal the composite nature of MMN, and suggest that breaking down MMN into distinct subcomponents may provide a more complete biomarker for pathological conditions associated with unusual prediction mechanisms.

## 2. Methods

### 2.1. Participants

Thirty participants were recruited in this study (15 males and 15 females, age: 24 ±2.6 years old, mean ± standard deviation). The participants self-reported and were screened to have no participation in drug studies, no serious deficits in vision and hearing, and no history of neurological and psychological conditions. This study was approved by the Research Ethics Committee of the National Taiwan University Hospital (201906081 RINA), and all participants gave written informed consent after understanding experimental procedures and before the experiment.

### 2.2. Local-global paradigm with controlled predictabilities

We implemented an auditory local-global oddball paradigm, creating 8 distinct sequence blocks (Blocks 1 to 8). Each block features a unique combination of local and global regularities, formed through varying sequence lengths and occurrences (Figure 1). Two different tones were created by combining three sinusoidal waves of base frequencies: the low-pitched tone with 350, 700, and 1400 Hz, and the high-pitched tone with 500, 1000, and 1500 Hz. Each tone has a 100 msec duration with a 7 msec rise and fall. For each block, we delivered 144 tone sequences, each consisted of 2 or 3 tones consecutively delivered with a 200 msec stimulus onset asynchrony (SOA). The inter-trial interval (ITI) between the end of one sequence’s last tone and the start of the next sequence’s first tone was randomly set to a value between 1000-1400 msec, in 50 msec increments.

**Figure 1.**
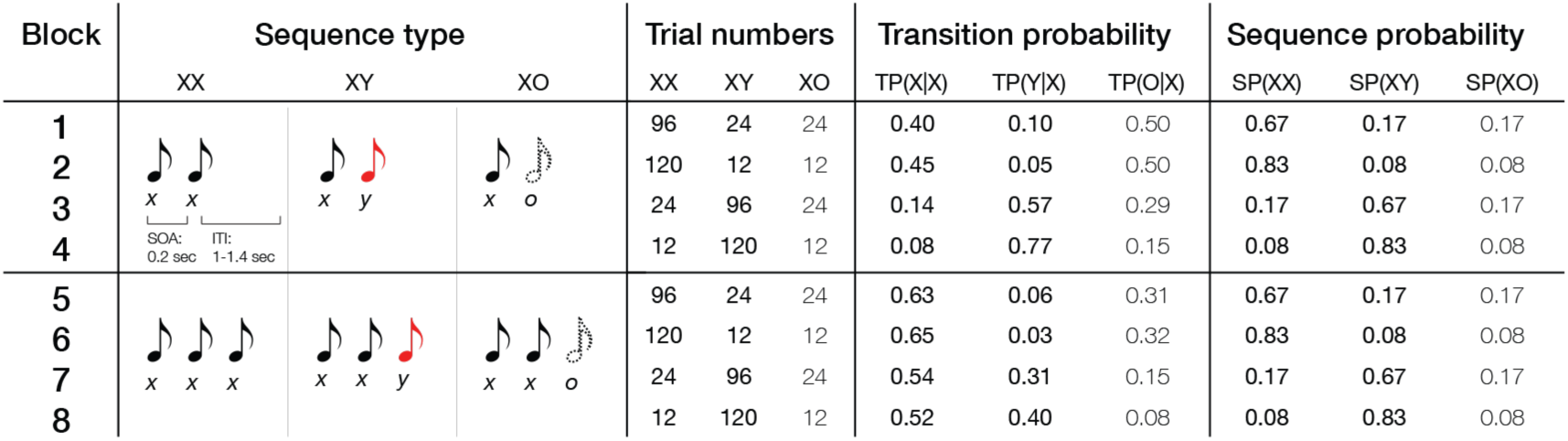
Task design. The configuration of sequence types, number of trials, transition probabilities, and sequence probabilities in 8 blocks. The tone icons colored in black and red represent tone x and y in different pitches while the tone icon with the dashed outline represents omission (“o”, no tone delivered). Sequences *xx* and *xxx* belong to Sequence type XX. Sequences *xy* and *xxy* belong to Sequence type XY. Sequences *xo* and *xxo* belong to Sequence type XO. The probabilities were rounded to two decimal places.

In Blocks 1 to 4, there were three types of tone sequence: *xx* represented two tones with the same tone pitch, *xy* represented that the last tone differed from the preceding tone, and *xo* represented that the last tone was omitted. Blocks 5 to 8 contained 3-tone sequences: *xxx*, *xxy*, or *xxo*. Note that *xx* and *xxo* are identical sequences, but the latter represents an omission in the blocks of 3-tone sequences. Collectively, the sequences *xx* and *xxx* are labeled as sequence type XX, while *xy* and *xxy* are labeled as XY, and *xo* and *xxo* are labeled to as XO.

The numbers of trials for sequence types XX, XY, and XO were set to be 96:24:24 for Blocks 1 and 5, 120:12:12 for Blocks 2 and 6, 24:96:24 for Blocks 3 and 7, and 12:120:12 for Blocks 4 and 8. The order of the 144 sequences in each block was pseudorandom. Each block was sectioned into four phases in which the proportion of sequence types was maintained (e.g., in Block 1, every phase contained 24 trials of XX, 6 trials of XY, and 6 trials of XO). Within each phase, the order of the 36 sequences was random. The 8 blocks were delivered twice: once with the low-pitched tone as tone *x* and the high-pitched tone as tone *y*, once with the high-pitched tone as tone *x* and the low-pitched tone as tone *y*.

With different sequence lengths and ratios, each block had a unique configuration of local and global regularities. The local regularity was established by the tone-to-tone transition probability (TP), i.e., the conditional probability of an incoming tone given the previous tone. TP in each block contained three values: TP(X|X), TP(Y|X), and TP(O|X), which represent the conditional probabilities of tones *x*, *y*, and o occurring, given the previous tone *x*, within a single block. On the other hand, the global regularity was established by the multi-tone sequence probability (SP). SP in each block contained three values: SP(XX), SP(XY), and SP(XO), which represent the probabilities of sequence types XX, XY, and XO, within a single block. The 8 blocks featured distinct combinations of TP and SP (Figure 1), which allowed us to control the predictability of sensory stimuli and examine varying degrees of prediction error at both local and global levels. The TP and SP calculations are shown in Supplementary Method.

During the experiment, participants were instructed to pay attention to the sound while visually fixating at a central fixation (white cross), and no behavioral response was required. A task-irrelevant video was displayed during the break between two block presentations (runs) to minimize the influence of the learned regularities in a run being carried over to the next run. All stimulus presentation were programmed with MATLAB-based PsychtoolBox (Kleiner et al., 2007; Pelli, 1997) and presented with a monitor (resolution: 1920*1220 pixels; sampling rate: 60 Hz) and a pair of desktop speakers (~60 dB).

### 2.3. EEG recording and analysis

Raw EEG data was recorded with a 64-channel QuickCap (Compumedics NeuroScan Inc.). During recording, the data was referenced to a reference electrode near Cz, and impedances were kept less than 2 kΩ for two mastoid electrodes and 5 kΩ for the remaining electrodes. We set a band-pass filter of 0.01 to 100 Hz and a 500-Hz sampling rate.

We used MATLAB-based EEGLAB (Swartz Center for Computational Neuroscience)(Delorme & Makeig, 2004) to preprocess the data from each participant. First, we merged the data from all runs and re-referenced it to the average of two mastoid electrodes to eliminate systematic noise from the environment (functions: *pop_mergeset.m*; *pop_reref.m*). Second, epochs of sequences were extracted from 1.2 sec before the first tone to 1.9 sec after the last tone (*pop_epoch.m*). Third, bad epochs containing excessive fluctuations or high-frequency noise were manually removed. On average, approximately 2% of the total epochs in each participant were removed at this step. Fourth, we removed eye and muscular artifacts using the ADJUST toolbox (An Automatic EEG artifact Detector based on the Joint Use of Spatial and Temporal features) (Mognon et al., 2011) (*pop_runica.m*, *interface_ADJ.m*, *pop_subcomp.m*). Finally, the processed epochs were corrected with the baseline estimated within the time window of –300 to –100 msec relative to the onset of the first tone (*pop_rmbase.m*). We down-sampled the data to 250 Hz to speed up later analyses. For each participant, channel, and block, we then averaged the signals across trials to obtain the event-related potential (ERP) for each sequence type (i.e., XX, XY, and XO). To measure deviant responses, we then compared ERPs from the sequence type XY to ERPs from the sequence type XX (XY – XX) for each participant, channel, and block. Examples of the deviant responses from – 200 to 800 msec relative to the onset of the last tone are shown in Figure 2.

**Figure 2.**
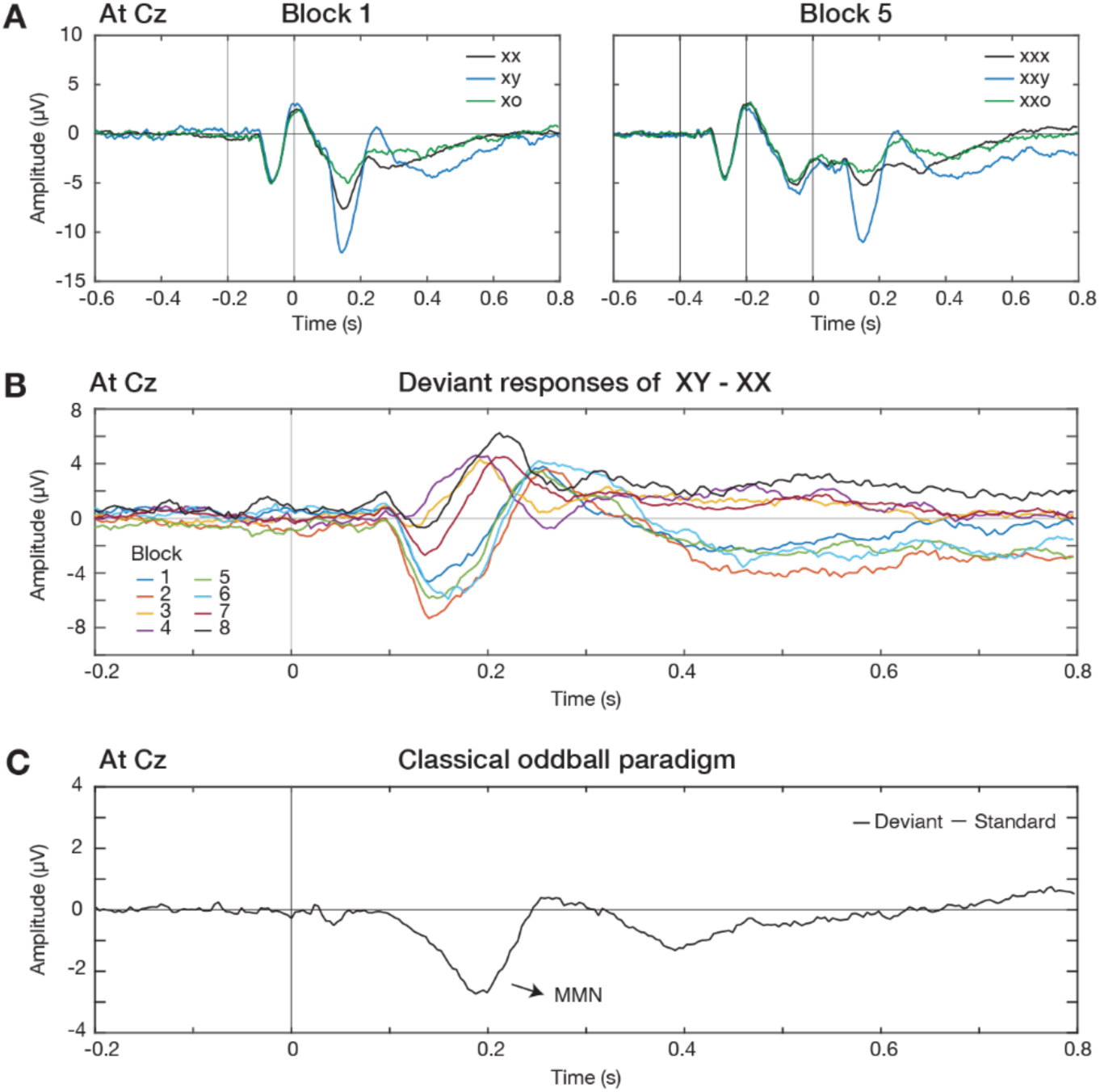
ERPs and deviant responses from contrasts. (A) The ERPs of sequences *xx*, *xy*, and *xo* in Blocks 1 (left panel) and sequences *xxx*, *xxy*, and *xxo* in Block 5 (right panel) at channel Cz. Solid vertical lines represented the stimuli of a sequence, and time zero was set to be the onset of the last stimulus. (B) The deviant responses obtained by contrasting ERPs of sequence types XY and XX in 8 blocks. (C) An example of MMN response in the classical oddball paradigm (open data from Kappenman et al., 2021)

### 2.4. Regression analysis

To examine the effects of the local regularity (TP) and the global regularity (SP) on the MMN, we implemented linear mixed-effect model analysis (function: *lmer*), using R (version 4.2.2). To select the peak amplitude and peak latency of the MMN for each participant and block, we focused on the deviant responses at Cz and in the range of 100-250 msec after the onset of the last tone (Näätänen et al., 2007) (Supplementary Figure 1). Four models were created: (1) TP only, (2) SP only, (3) both TP and SP, and (4) both TP and SP with an interaction term (TP*SP) as the fixed effect (see Table 1). The participants were set as the random effect of intercept.

**Table 1.**
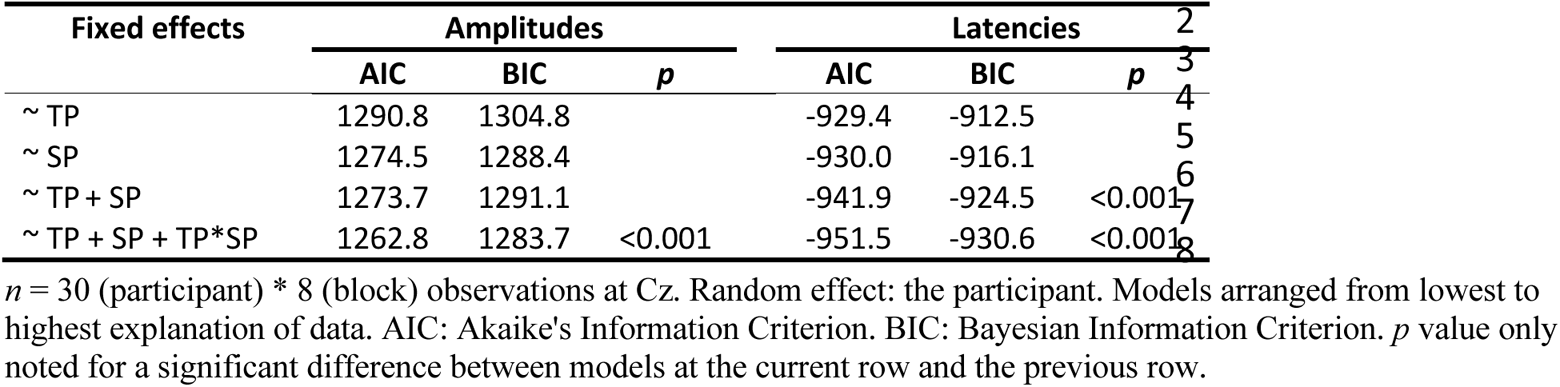
Comparisons of linear mixed-effect models on the MMN.

Since it has been found that MMN changes with occurrences of novel stimuli (Sato et al., 2000), we used TP(Y|X) (i.e., the transition probability from tone *x* to tone *y*) to represent TP, and SP(XY) (i.e., the sequence probability of sequence type XY) to represent SP. The goodness of fit in models penalized by model complexity was evaluated with Akaike’s Information Criterion (AIC) and Bayesian Information Criterion (BIC), and alpha was set to be 0.05 (R function: *anova*).

### 2.5. Parallel factor analysis

To reveal underlying subcomponents in deviant responses, we implemented the parallel factor analysis (PARAFAC) (Harshman & Lundy, 1994), a generalization of principal component analysis to higher orders. We first pooled the total deviant responses into a 3-dimensional tensor (62 channels × 250 time points × 8 blocks), and used PARAFAC to decompose the tensor into multiple subcomponents. The analysis was implemented by using the N-way toolbox (Andersson & Bro, 2000) (function: *parafac.m*). The convergence criterion was set to be 1e-6, and the three dimensions were not constrained in terms of orthogonality or positivity.

PARAFAC was performed with varying numbers of subcomponents, ranging from 1 to 8. To determine the optimal number of subcomponents in the tensor, we evaluated the Core Consistency Diagnostic (CORCONDIA) (Bro & Kiers, 2003; Harshman & Lundy, 1994). Decomposition with low consistency indicates a poor appropriateness, where high interactions exist between subcomponents. A CORCONDIA of 80~90% is considered a good decomposition and below 50% considered a problematic decomposition (Bro & Kiers, 2003; Pouryazdian et al., 2016). Besides CORCONDIA, we also evaluated BIC to quantify how well the data decomposition fit the observed data. The optimal number of subcomponents was determined by high CORCONDIA and low BIC.

Two subcomponents (Subcomponents 1 and 2) were identified from PARAFAC (see details in Results). The spatial, temporal, and functional characteristics of each subcomponent can be described by the score or loading matrix (activation values) in its original dimensions: *Channel*, *Timecourse*, and *Block*.

### 2.6. Functional roles of subcomponents identified by a predictive coding model

To verify the functional roles of the identified subcomponents, we used a predictive coding model which can estimate the size of prediction errors during the local-global oddball paradigm (Chao et al., 2022). The model quantitatively describes prediction and prediction-error signals at two hierarchical levels, where the first and second levels encode predictions based on TP and SP, respectively. Using the formula and codes provided by Chao et al. (2022), we calculated the theoretical quantities for both the local (or first-level) prediction-error signal and the global (or second-level) prediction-error signal within the deviant responses. The local prediction-error signals across the 8 blocks were represented by 8 model values (*PE_1_*), while the global prediction-error signals across the same blocks were characterized by 8 model values (*PE_2_*).

To examine whether Subcomponents 1 and 2 were associated with the local and global prediction-error signals, we first quantified the Pearson’s correlation coefficients between their 8 activation values in the *Block* dimension (denoted as *Block_1_* for Subcomponent 1 and *Block_2_* for Subcomponent 2) and the model values (*PE_1_* and *PE_2_*). We also reconstructed deviant responses from Subcomponents 1 and 2 with *PE_1_* and *PE_2_* and compared them to the actual responses. The response at time *t* for channel *i* and block *j* was reconstructed (denoted as *Recon*) as follow:

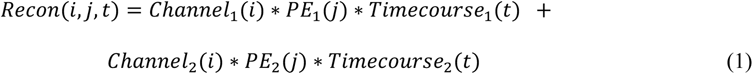

where *Channel_1_* represents the 62 activation values in the *Channel* dimension for Subcomponent 1, and *Timecourse_1_* represents the 250 activation values in the *Timecourse* dimension for Subcomponent 1. Similarly, *Channel_2_* and *Timecourse_2_* represent those for Subcomponent 2.

The reconstruction with the opposite association, i.e., Subcomponent 1 as *PE_2_* and Subcomponent 2 as *PE_1_*, was also tested:

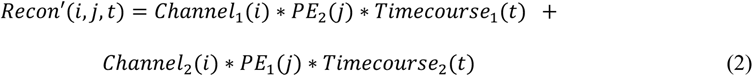

We then calculated the mean squared difference, *MSD* and *MSD’*, between reconstructed responses and the actual response, *Act*, for each channel *i* and block *j*:

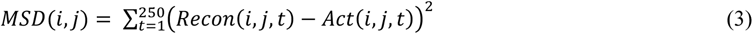

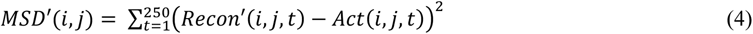

We also measured the Pearson’s correlation coefficients of the reconstructed and actual time courses (*t* = 1:250) for each channel *i* and block *j*:

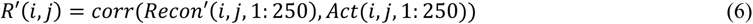

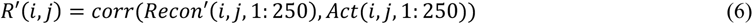

To evaluate which association produced better reconstruction, *MSD* and *MSD’* were compared across 62 channels and 8 blocks using paired t-test (R function: t_test) and Cohen’s D for effect size (R function: cohens_d). Similarly, *R* and *R’* were compared across 62 channels and 8 blocks.

## 3. Results

### 3.1. Deviant responses varied across blocks

Figure 2A shows an example of ERPs, averaged across participants, of the three sequence types from the channel Cz in Blocks 1 and 5. For each block, we extracted the deviant responses by contrasting ERPs between the sequence type XY and XX (XY *–* XX). Figure 2B shows the average deviant responses from the 8 blocks. The deviant responses were found to vary across blocks, particularly around the typical timing of MMN (100 to 200 msec) and around the typical timing of P300 (200 to 300 msec). For comparison, Figure 2C shows an example of MMN at the same channel from published data of a classical auditory oddball paradigm where the probability of the deviant tone was 20% (Kappenman et al., 2021). The MMN from the classical oddball paradigm shared a similar negative waveform with some of the deviant responses in our local-global oddball paradigm.

We also estimated the omission error by contrasting ERPs between the sequence type XO and XX (XO *–* XX) (see Supplementary Figure 2). Positive responses were found around 100-200 msec. However, such contrast contains not only the omission error but also the sensory input of the last tone in the sequence XX. Because of this contamination, we did not include the responses of XO – XX for further analysis.

### 3.2. MMN amplitudes modulated by local and global regularities

To examine whether and how the local and global regularities (i.e., TP and SP) affect the amplitude of MMN, we regressed the peak amplitudes of MMNs in all blocks and participants (Supplementary Figure 1) against different probabilities. Among the four models (see Table 1 left column), the model that included TP, SP, and their interaction term had the smallest values of AIC and BIC. In this optimal model (Table 2, left column), there were a significant interaction term (*p* <0.001) and main effects of TP (*p* <0.001) and SP (*p* <0.001). The significant interaction term showed that the effect of TP on MMN was not independent but modulated by SP; on the other hand, the effect of SP on MMN was modulated by TP.

**Table 2.** Fixed effects on the MMN.

|  | Amplitudes |  |  | Latencies |  |  |
| --- | --- | --- | --- | --- | --- | --- |
|  | Estimates | SE | *p* | Estimate<br>s | SE | *p* |
| (Intercept) | -10.1 | 0.6 | <0.001 | 0.18 | 0.01 | <0.001 |
| TP | 21.9 | 5.5 | <0.001 | -0.12 | 0.06 | 0.04 |
| SP | 7.5 | 1.4 | <0.001 | -0.08 | 0.02 | <0.001 |
| TP*SP | -24.0 | 6.6 | <0.001 | 0.24 | 0.07 | <0.001 |

14 Estimate: correlation coefficient. SE: standard error.

### 3.3. MMN latencies modulated by local and global regularities

We also estimated the effects of the local and global regularities on the temporal characteristic of MMN: peak latency. Table 1, right column, shows that the peak latencies were also best explained by the model including TP, SP, and their interaction term. There were significant interaction (*p* <0.001) and significant main effects of TP (*p* = 0.04) and SP (*p* <0.001) (Table 2, right column). The significant interaction indicated that the interaction between local and global regularities was also shown in the MMN latencies.

### 3.4. Extracting MMN subcomponents

The linear mixed effect model analysis revealed that MMN was modulated by both TP and SP. This suggested that MMN could contain two overlapping subcomponents that respectively encode TP and SP. To test this, we first used PARAFAC to analyze the total deviant responses, aiming to identify the number of latent subcomponents contained within them.

We decomposed the total data with 1 to 8 subcomponents and quantified the CORCONDIA for each decomposition. We found an abrupt drop in CORCONDIA, from nearly 100% to 18%, when the decomposition changed from 2 subcomponents to 3 subcomponents (Supplementary Figure 3). This indicated that the optimal number of subcomponents in the deviant responses was two. Furthermore, a lower BIC value was found in two subcomponents, also indicating that the responses can be better explained by the two subcomponents (−0.76e+05) than one subcomponent (−0.30e+0.5).

The two subcomponents, Subcomponents 1 and 2, are visualized in the dimensions of space (*Channel*), time (*Timecourse*), and function (*Block*) (see Figure 3). Spatially, Subcomponent 1 was found in the central-frontal area, while Subcomponent 2 was found in a more frontal area (Figure 3A). Temporally, both components showed negative waveforms after the last tone but with a delay between them (Figure 3B). Subcomponent 1 peaked negatively at 136 msec, followed by a positive response during 200-400 msec. Subcomponent 2 peaked negatively at 196 msec, followed by a long-lasting negative response. Functionally, the two subcomponents had different contributions to the deviant responses across the 8 blocks (Figure 3C, blue line).

**Figure 3.**
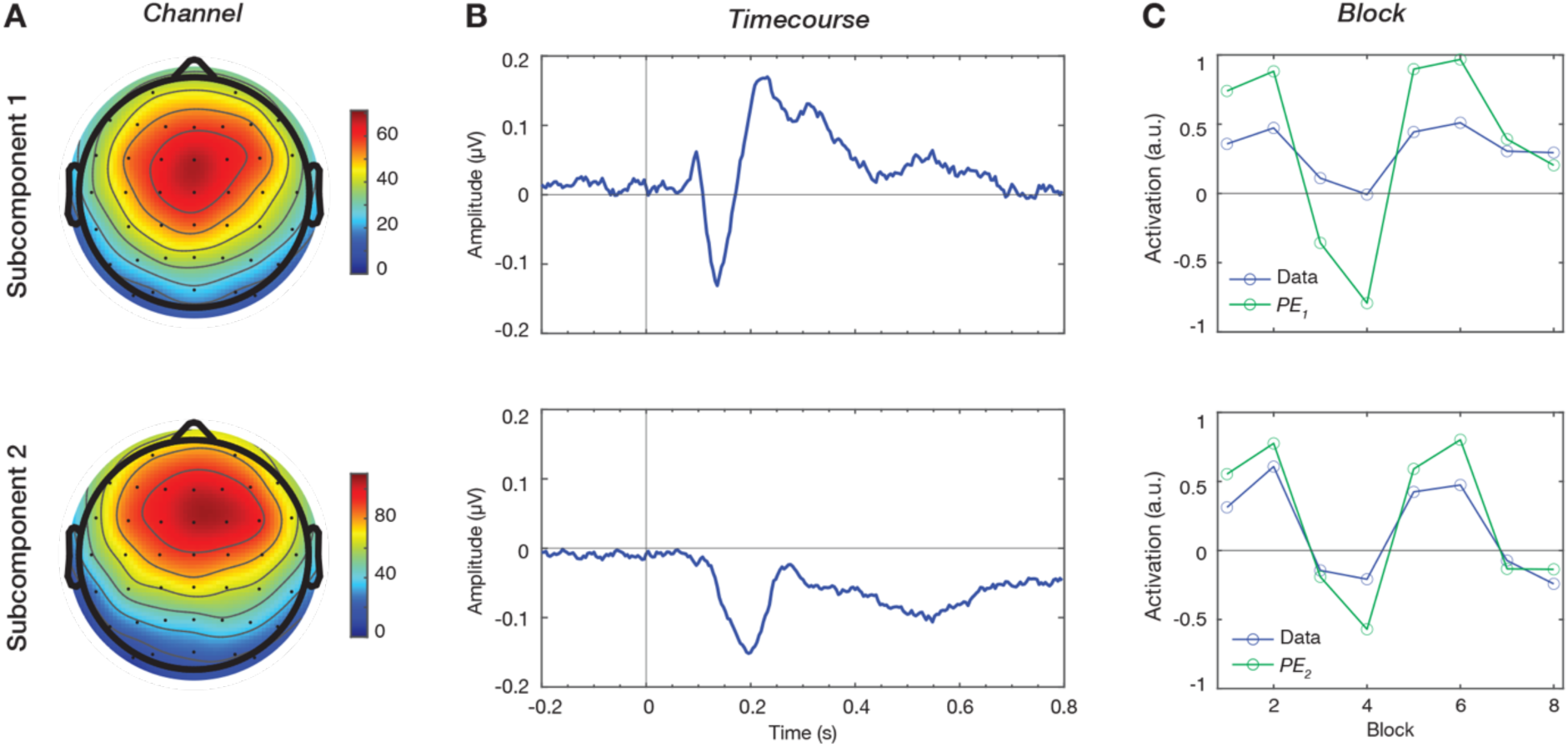
Neural signatures of prediction-error signals. Subcomponents 1 and 2 obtained from the data-driven hypothesis-free decomposition on 8 deviant responses. The two extracted components shown in the (A) *Channel*, (B) *Timecourse*, and (C) *Block* dimensions. The *Block* dimension represented activation of the components to the deviant responses (the blue line) and the model values of the first- and second-level prediction errors, denoted as *PE_1_* and *PE_2_* (the green line).

### 3.5. Functional roles of the MMN subcomponents

To identify functional roles of Subcomponents 1 and 2, we compared their activation profiles across the 8 blocks (*Block_1_* and *Block_2_*) to the theoretical values of the local and global prediction errors (*PE_1_* and *PE_2_*) obtained from a hierarchical predictive coding model (Figure 3C, green line).

Based on their spatiotemporal characteristics, we hypothesized that Subcomponent 1, occurring earlier in a less frontal area, represented the local prediction error. On the other hand, Subcomponent 2, occurring later in a more frontal area, was hypothesized to represent the global prediction error. This hypothesis was first supported by the high correlation coefficients between *Block_1_* and *PE_1_* (r = 0.99, Pearson’s correlation coefficient) and between *Block_2_* and *PE_2_* (r = 0.96). To further validate the hypothesis, we reconstructed signals with the model values and compared them to actual deviant responses.

Figure 4A showed examples of reconstructed time courses at channel Cz in Block 7. In this example, *Recon*, which associated Subcomponent 1 with *PE_1_* and Subcomponent 2 with *PE_2_*, exhibited a response similar to the actual one. In contrast, *Recon’*, with the reversed association, exhibited a response that was markedly different from the actual one. Across all channels and blocks, *MSD* was significantly smaller than *MSD’* (t[495] = −12.05, *p* = <0.001, d = −0.54 [moderate]) (see Figure 4A). Furthermore, *R* was significantly higher than *R’* (t[495] = −8.27, *p* = <0.001, d = −0.37 [small]) (see Figure 4B). These results suggested that Subcomponents 1 and 2 represented the local and global prediction-error signals, respectively.

**Figure 4.**
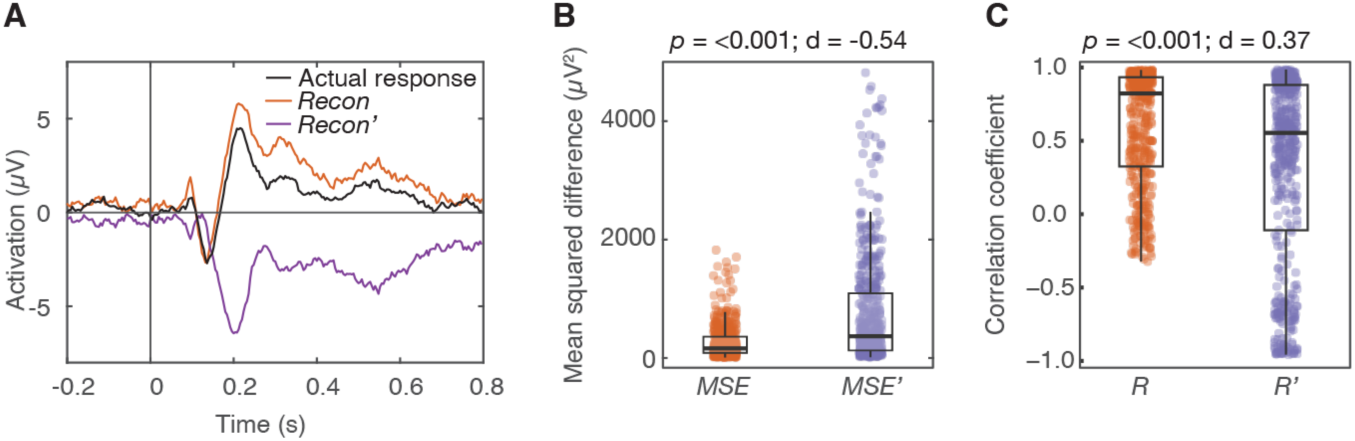
Reconstruction with functional assignments and their differences from the actual ERP. (A) Examples of reconstructed signals at Cz in Block 7. The black line represents the actual deviant response. The orange line represents Recon, the reconstructed deviant response with Subcomponent 1 as *PE_1_* and Subcomponent 2 as *PE_2_*, while the purple line represents *Recon*’, the reconstructed response with the opposite assignment. Mean squared differences in the panel B and Correlation coefficients in the panel C were calculated across channels and blocks (62*8 dots for each assignment). *p* value was obtained using paired T-test and d value was Cohen’s D.

## 4. Discussion

In the current study, we combined an extended auditory local-global oddball paradigm with a data-driven decomposition analysis to extract subcomponents in MMN. We further demonstrated that the early MMN subcomponent in the central-frontal area represents the local prediction-error signal while the late MMN subcomponent in the more frontal area represents the global prediction-error signal. Our study applies the predictive-coding framework to reveal the complex nature of MMN, and establishes a robust experimental and analytical platform that can examine the functionality of multilevel deviant detection in both healthy and affected brains.

### 4.1. Hierarchical MMN in predictive-coding framework

We revealed that the MMN of the deviant response varied with the hierarchical probabilities of the stimulus presentation and notably consisted of two MMN subcomponents with distinct spatial and temporal characteristics obtained from an unbiased decomposition.

The two subcomponents spatially complied with the hierarchical network which is the 3-level hierarchy of cortical cascades in responses to auditory changes consisting of bilateral primary auditory cortices receiving inputs, superior temporal gyrus functioning as memory trace of priors, and right inferior frontal gyrus modulating the attention allocation (Doeller et al., 2003; Giard et al., 1990; Halgren et al., 2011; Hofmann-Shen et al., 2020; MacLean & Ward, 2014; Opitz et al., 2002). This was also modeled in different types of oddball paradigms (Garrido et al., 2007, 2008). In our findings, the early MMN was activated in the frontal-central area, which is consistent with previous literature and may reflect dipole sources in the superior temporal area (Alho et al., 1998; Molholm et al., 2005; Schönwiesner et al., 2007). On the other hand, the late MMN was activated in the more frontal area and slightly dispersed toward the right side, which may be mapped to an anterior portion of the brain such as the right prefrontal cortex. Although EEG signals are limited in spatial resolution, the temporal patterns of the EEG signals may provide some hints regarding potential sources. Specifically, we found that the early subcomponent (i.e., Subcomponent 1) of MMN at the central-frontal area was elicited earlier and followed by the late subcomponent (Subcomponent 2) of MMN in the frontal area, which is consistent with the differences in the peak latency found in the temporal and frontal generations of MMN (Rinne et al., 2000) and bottom-up gamma oscillations (Chao et al., 2022). Furthermore, in alignment with the hierarchical predictive coding theory, the observed temporal pattern offers directional support for feedforward error propagation between low and high cortical hierarchy, and activations of MMN subcomponents closely resembled the quantitative definition of local and global prediction-error signals (Chao et al., 2022).

However, the findings and interpretation of the early and late subcomponents of MMN appear to deviate from previous studies of the local-global oddball paradigm claiming that the MMN represents the local prediction-error signal while the P300 represents the global prediction-error signal (Wacongne et al., 2011). We argued that this discrepancy results from the difficulty of dissociating hierarchical prediction-error signals. P300 in previous literature was typically captured by comparing responses to stimuli of (global deviant + local deviant) + (global deviant + local standard) to stimuli of (global standard + local deviant) + (global standard + local standard). It was assumed that such contrast would extract the deviance between the global deviant and the global standard, while the deviance at the local level was assumed to be controlled. In fact, however, the local prediction-error signal cannot be fully canceled out between blocks because the blocks had different layouts of the stimulus occurrence, leading to different transition probabilities and thus different local predictability. In contrast, our approach adopted a decomposition approach to extract subcomponents from ERP contrasts which were correlated with both the local and global prediction-error signals. As the reconstruction of two components showed, the late positive waveform was the product of the subcomponent following the early MMN (Alho et al., 1998; Näätänen et al., 2005), and the inverse late MMN representing the negative prediction error at the global level. Moreover, the late MMN peaking at about 200 msec shared similarities with the N2b component, a negative ERP in the N2 family occurring around 200 to 300 msec and after MMN (also known as N2a) (Folstein & Van Petten, 2008; Näätänen & Picton, 1986; Pritchard et al., 1991). It has been studied in domains of attention, response inhibition, and cognitive control, where participants are required to pay active attention or make behavioral responses (Barry & De Blasio, 2015; Kasai et al., 1999). This relatively late-stage process coincides with cognitive demands for attentional control and consciousness in learning the global regularity in the local-global task (Bekinschtein et al., 2009).

### 4.2. Potential for clinical application

The hierarchical MMN discovered in the local-global oddball paradigm provides a platform to link atypical deviant responses across multiple levels to disease heterogeneity. For instance, individuals with ASD have been found to have overall smaller MMN responses compared to their typically developing controls (Lassen et al., 2022; Vlaskamp et al., 2017). Also, differences in MMN between the predictable and unpredictable sequence structures are smaller in ASD, suggesting less modulation by global regularity (Goris et al., 2018). With the decomposition approach in the current study, activations of the early and late subcomponents differing from healthy individuals could reflect inefficiencies in learning local and/or global regularities to detect deviants. For example, smaller differences in MMN between the predictable and unpredictable sequence structures could result from low activation of the late MMN subcomponents or low activations of both the early and late MMN subcomponents. Such atypical activations of the subcomponents may also differ across individuals with ASD and reflect characteristics of the heterogeneous nature of the disorder. Taking a broader view, this approach could also be applied to help delineate other mental illnesses or developmental delays exhibiting difficulties in detecting deviants embedded in multi-level statistical regularities.

To increase the feasibility of the local-global oddball paradigm in clinical settings, we tested the robustness of implementing the decomposition analysis on the data from a short recording protocol which was similar to the classical local-global oddball paradigm, and then used the decomposition analysis. The optimal number of the extracted components from the decomposition was 2 (Supplementary Figure 4), and the subcomponents replicated the results in Figure 3 (Supplementary Figure 5). With the advantages of a short and undemanding experimental protocol, future research could continue to test its reliability and link to symptoms in clinical populations.

### 4.3. Limitations and future research

Although our findings provide a more thorough understanding of hierarchical MMN, there are still some limitations to be resolved and advances to be achieved in future research: hierarchical structure composed of more than two levels, prediction updates over time, and neuronal mechanism of learning statistical regularities at two levels.

Firstly, the functional hierarchy underlying MMN is not limited to two levels, as the prediction error likely does not propagate only across two levels of the hierarchical scheme. For example, the encoding of natural images was modeled with the predictive coding perspective and multiple layers simulating visual cortical processing (Rao & Ballard, 1999; Spratling, 2017), and temporal sequences can consist of not only single-stimulus transition rules but also arbitrary regularities (Dehaene et al., 2015). In order to extract multi-level prediction-error signals in a real sequential environment, probability calculations for different regularities and quantitative values of the prediction error at each level should be determined.

Secondly, the prediction error is generated to update the precision of the prediction; that is, a prediction update follows a prediction error evoked by sensory input. In this study, we observed that the global prediction-error signals had a long-lasting negative waveform after MMN, which might be related to desynchronized beta oscillations as prediction updates (Bastos et al., 2012; Chao et al., 2018). However, our study is incapable of dissociating the prediction update and the prediction error because of their interdependence and of evaluating the dynamics of prediction updates during learning. Future research could incorporate analytical technique with focus on temporal order to dissociate the prediction update from the prediction error, and trial-by-trial analysis to capture the dynamics.

Thirdly, for the oddball paradigm with the complex sequence structure, the underlying mechanism of how multiple regularities are computed at the neuronal level and the links between the neuronal computations of the regularities and the MMN subcomponents are critically needed for investigation. We could consider a combination of a biologically realistic model and a microscopical approach such as calcium imaging and high-density neural recording in the future.

## Data availability

The raw EEG data was originally used in Chao et al. (2022) and can be obtained from the corresponding author upon request.

## Contributions

Z.C.C. and Y.T.H conceptualized the study. Y.T.H, C.W., and Z.C.C refined the experimental protocol. Y.T.H collected the data and performed data analysis. Y.T.H wrote the draft of the paper, and Z.C.C, C.W and S.K. helped with editing. All authors contributed to and approved the final paper.

## Declaration of competing interest

There is no conflict of interest related to this work for any of the authors.

## Supporting information

all_materials

## Acknowledgments

We thank Hsuan-Chi Liu, Chin-Kun Fu, and Shih-Yao Mao for helping with participant recruitment and experiment preparation. We also thank Felix B. Kern and Amit Yaron for proofreading. This study was supported by Japan Society for the Promotion of Science, Japan (to Y.T.H), Ministry of Science and Technology of Taiwan, Taiwan (MOST 106-2420-H-002-008-MY2 and MOST 109-2410-H-002-106-MY3) (to C.W), and World Premier International Research Center Initiative (WPI), MEXT, Japan (to Z.C.C.).

## Notes

### Competing Interest Statement

The authors have declared no competing interest.

