## Supplementary material for "Dissecting Mismatch Negativity: Early and Late Subcomponents for Detecting Deviants in Local and Global Sequence Regularities": all_materials

**Title**

***Correspondance:**

Zenas C. Chao

Key terms:

Mismatch negativity, hierarchy, subcomponents, predictive coding, EEG

**Supplementary Method**

**Calculations of transition probability (TP) and sequence probability (SP)**

The transition probability (TP) in each block contained three values: TP(X|X), TP(Y|X), and TP(O|X) for each block, which represent the conditional probabilities of tone x, y, and o given the previous tone x, respectively. For *n*-tone sequences, there are *n*-1 transitions from *x* to *x* (*TNx*) and 1 transition from *x* to *o* (*TNo*) in the sequence type XX. In the sequence type XY, there are *n*-2 *TNx* and 1 transition from *x* to *y* (*TNy*). In the sequence type XO, there are *n*-2 *TNx* and 1 *TNo.* Note that the *x*-to-*o* transition was considered at the end of the tone sequence *xx*, since the sequence ending cannot be known by stimulus transitions but rather by sequence structure (Chao et al., 2022). Thus, the transition from tone *x* continues to predict the incoming input (tones *x*, *y*, or *o*). Then, based on the trial numbers of the sequence types, we calculated all instances of the three transition types within one block, and the corresponding transition probabilities. Taking 2-tone sequences in Block 1 as an example, there were (1 *TNx* and 1 *TNo*)*96 in total in 96 presentations of tone sequence *xx.* There were 1 *TNy**24 in 24 presentations of tone sequence *xy*. There were 1 *TNo**24 in 24 presentations of tone sequence *xo*. Therefore, TP(X|X), TP(Y|X), and TP(O|X) are 0.4 (96/240), 0.1 (24/240), and 0.5 (120/240).

The sequence probability (SP). SP in each block contained three values: SP(XX), SP(XY), and SP(XO), which represented probabilities of sequence types XX, XY, and XO, respectively. Taking Block 1 as an example again, there were 96 presentations of tone sequence *xx*, 24 of tone sequence *xy*, and 24 of tone sequence *xo*. Thus SP(XX), SP(XY), and SP(XO) are 0.64 (96/144), 0.17 (24/144), and 0.17 (24/144), respectively.

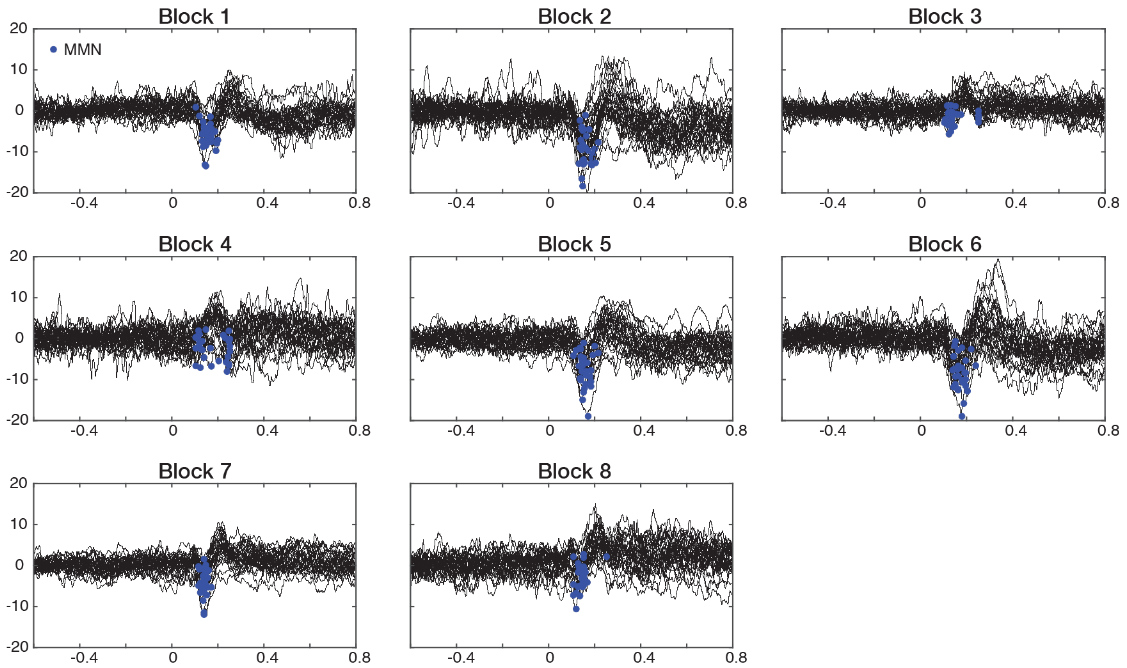


**Supplementary Figure 1. Deviant responses in 30 participants and their peak MMN.** Time zero represented the onset of the last stimulus. The blue dots (n = 30) represented peaks of MMN responses.

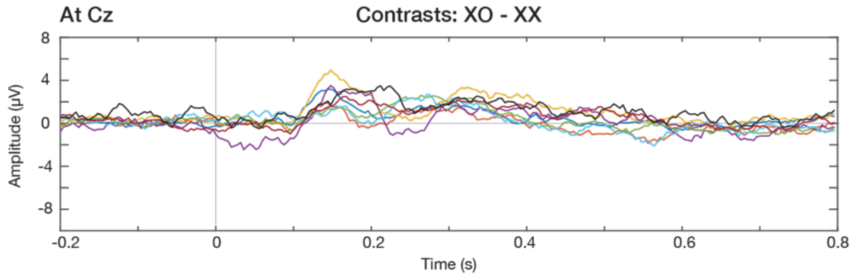


**Supplementary Figure 2.** Contrast responses by comparing ERPs of sequence types XO to XX.

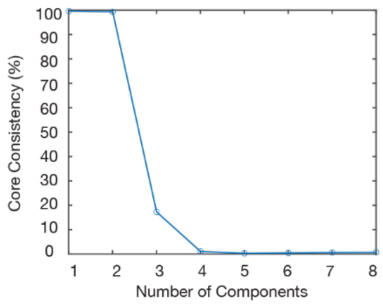


**Supplementary Figure 3. Core Consistency Diagnostic (CORCONDIA)**
The decomposition was conducted with 8 deviant responses.

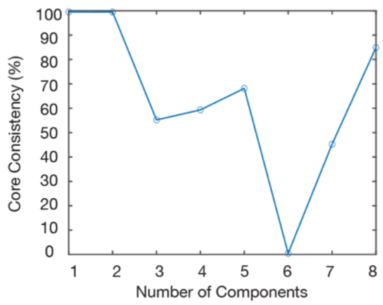


**Supplementary Figure 4. CORCONDIA**
The decomposition conducted with deviant responses in Blocks 6 and 8.

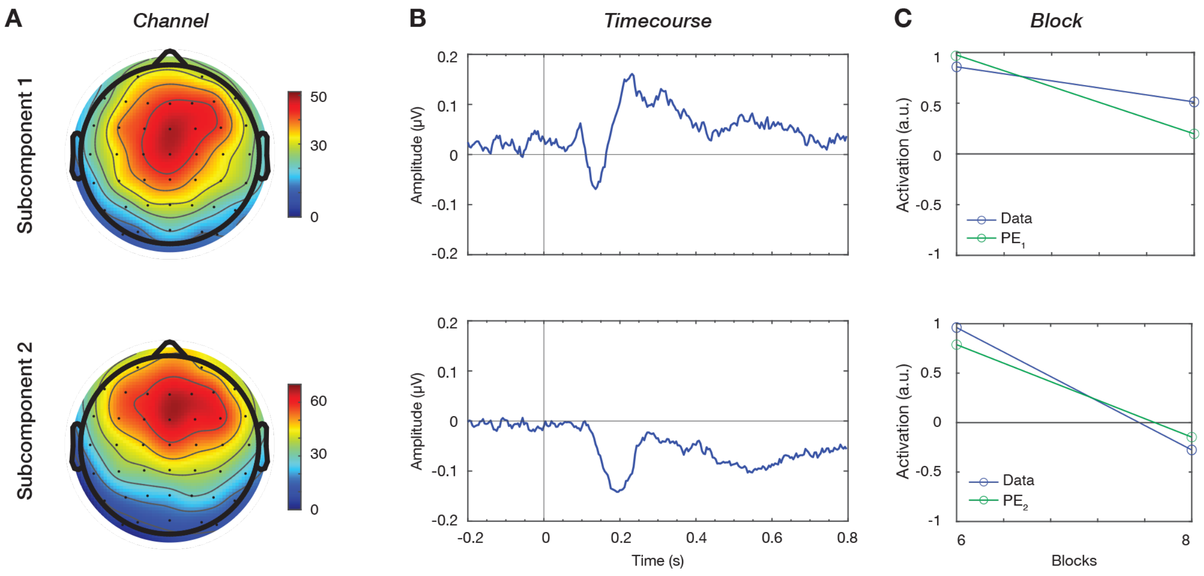


**Supplementary Figure 5. Extracted components.** Subcomponent 1 and 2 obtained from the decomposition on deviant responses in Blocks 6 and 8. The two extracted components shown in the (A) *Channel*, (B) *Time course*, and (C) *Block* dimensions.
